# Unexpected higher resilience to distraction during visual working memory in schizophrenia

**DOI:** 10.1101/567859

**Authors:** Yijie Zhao, Xuemei Ran, Li Zhang, Ruyuan Zhang, Yixuan Ku

**Affiliations:** The Shanghai Key Lab of Brain Functional Genomics, Shanghai Changning-ECNU Mental Health Center, School of Psychology and Cognitive Science, East China Normal University, Shanghai, 200062 China; Center for Magnetic Resonance Research, Department of Radiology and Neuroscience, University of Minnesota, Minneapolis, MN 55455 USA

**Keywords:** Schizophrenia, Visual working memory, distractibility, Bayesian observer model

## Abstract

Visual working memory (VWM) and distractibility are two core executive functions in human cognition. It has been suggested that schizophrenia (SZ) patients exhibit worse VWM performance and lower resilience to distraction compared with healthy control (HC) subjects. Previous studies, however, have largely investigated these two functions separately. It still remains unclear what are the mechanisms of the deficits, especially the interactions between the two cognitive domains. Here we modify the standard delay-estimation task in VWM and explicitly add distractors in the task so as to examine the two domains simultaneously. We find that SZ indeed exhibit worse performance compared with HC in almost all VWM load and distraction levels, a result consistent with most prior experimental findings. But adding distractors does not selectively impose larger impacts on SZ performance. Furthermore, unlike most previous studies that only focused on behavioral performance, we use the variable precision model to disentangle the distraction effect on different computational components of VWM (resources and resources allocation variability etc.). Surprisingly, adding distractors significantly elevates resources allocation variability—a parameter describing the heterogeneity of resource allocation across different targets—in HC but not in SZ. This counterintuitive result suggests that the internal VWM process in SZ is less interfered by the distractors. However, this unexpected higher resilience to distraction might be associated with less flexible cognitive control mechanisms. In sum, our work demonstrates that multiple cognitive functions might jointly contribute to dysfunctions in SZ and their interactions might manifest differently from merely summing their independent effects.

## INTRODUCTION

Visual working memory (VWM) is a central cognitive ability that provides temporary storage and manipulation of information^1,2^. VWM deficits have been widely documented in people with schizophrenia (SZ)^3–7^. But the underlying mechanisms still remain unclear. Existing theories propose impaired sensory processing at the encoding stage of working memory as one candidate mechanism of the behavioral deficits^8^. Indeed, our sensory systems are often confronted with an immense amount of information that greatly exceeds the processing capacity^9^. However, working memory capacity is known to be limited^10,11^. The capacity limitation necessities a selection process that prioritizes task-relevant information and filters out task-irrelevant ones in order to optimize performance. This is particularly important when salient distractors are present and interfering with the processing of targets. The interference induced by distractors, so-called “distractibility”, has been shown to link with several key cognitive functions, such as working memory^12^, endogenous and exogenous attention^13^, perceptual and value-based decision^14^, response inhibition^15^, cognitive control^16^. Moreover, atypical distractibility has been discovered in several psychiatry disorders, including ADHD^17^, autism^18^, depression^19^.

A sizable amount of literature has suggested the aberrant distractibility in SZ ^20–24^. One standard approach to study distractibility is to impose distractors in some cue-based attention tasks. However, most studies found no significant deficits in cue-based attention tasks in SZ^25,26^. One possibility is that the cues and instructions in those tasks were quite simple and 100% valid. Simple cues ease the tasks and require less attentional control. By contrast, if probed in high-demanding attention tasks, SZ exhibit deficits in suppressing salient distractors^27,28^. These findings suggest that the distractibility deficits in SZ exist and might be more prominent at the presence of highly salient distractors.

Recent advances in the basic science of VWM demonstrate that behavioral performance in VWM tasks is mediated by multiple factors^29^. It has long been proposed that SZ has lower memory capacity but intact memory precision compared with healthy control (HC) subjects^4,30^. This view has been proposed in the studies that use standard VWM tasks without distractors. It remains unclear whether SZ have deficits in VWM processing when confronted with distractors. From the computational perspective, distractors may reduce memory capacity and/or impair memory precision. Unfortunately, most previous studies on SZ have examined distractibility and VWM deficits separately. Few studies have attempted to combine them and investigate their interaction effect. It remains two unanswered questions: (1) whether SZ have distractibility deficits in VWM; (2) if yes, which VWM component(s) such distractibility deficits will influence.

In this study, we aimed to combine the classical distraction and VWM experimental paradigm to simultaneously the two functions in SZ. To do so, we modified a standard VWM task—color delay-estimation task. In the color delay-estimation task, subjects need to memorize the colors of all presented items and after a short delay reproduce the color of one cued item. In our modified version (Fig. 1), subjects were instructed to memorize only a subset of presented items (i.e., targets) and ignore other items (i.e., distractors). We independently manipulated the target size and the distractor size to control VWM loads and distraction levels. Moreover, we employed the Variable Precision (VP) model^31^ explicitly estimate three key aspects of VWM— the amount of resources at different target size level, the variability of resource assigned across items, and the variability induced by choice. Therefore, the VP model allows us to quantify the distraction effect in the computational process of VWM.

**Figure 1.**
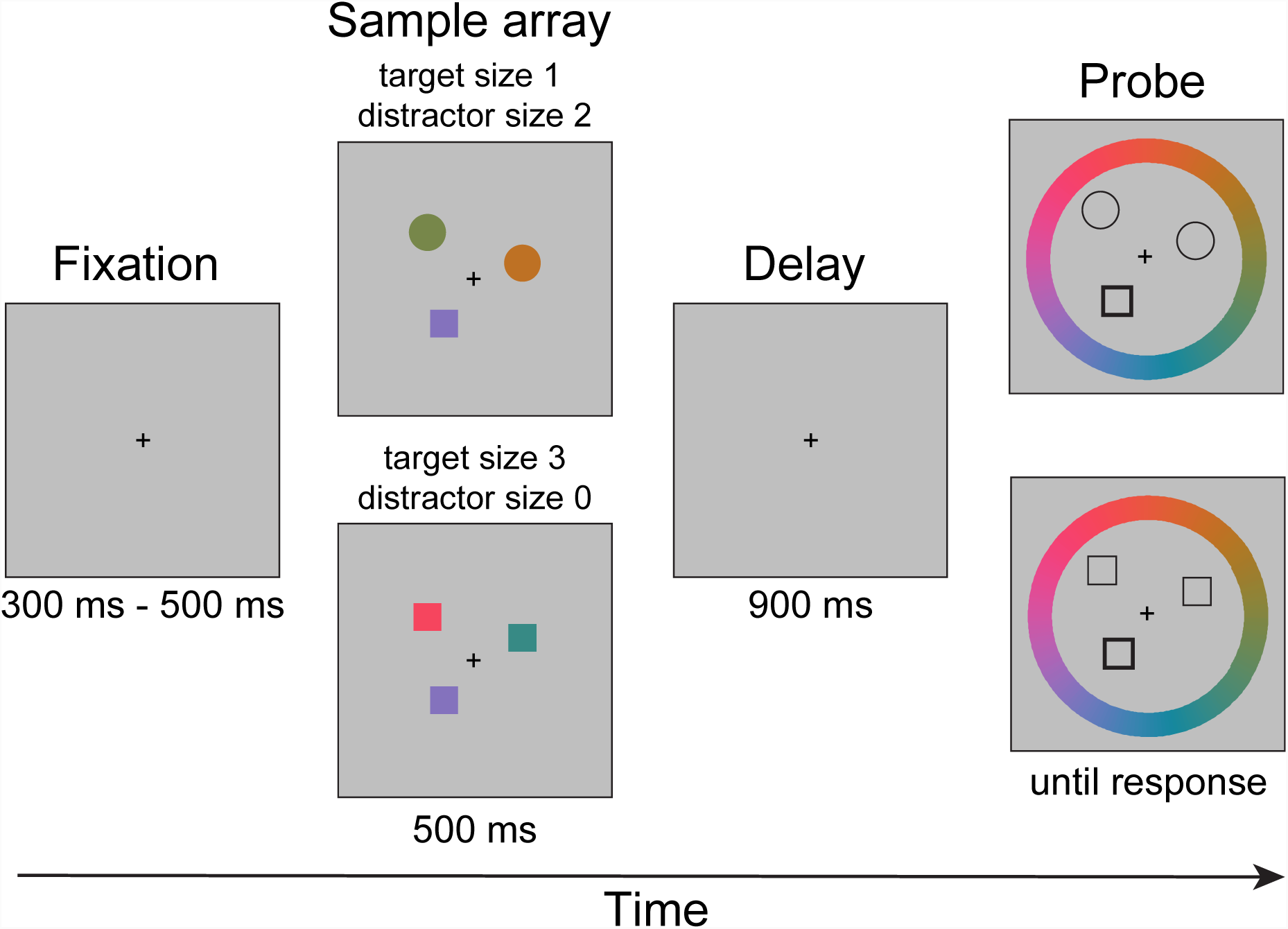
The modified color delay-estimation task. This figure illustrates two example trials of the experiment. In the experiment, each trial starts with a fixation point presented for 300ms to 500ms (with a step of 50ms). In the sample array, one or three targets (squares in this example) together with zero or two distractors (circles, a 2 × 2 design) are displayed on the screen for 500ms. Subjects were instructed to remember the colors of one of the shapes in the sample array. After a 900ms delay, outlines of the items at their original location would appear and one of the cued of target shapes is cued. Subjects are asked to recall and report the color of the target by clicking on the colored wheel using a computer mouse.

## MATERIALS AND METHODS

### Subjects

Sixty clinically stable SZ (33 inpatients and 27 outpatients) and sixty-one HC were recruited in this study. All SZ met the DSM-IV criteria for schizophrenia and were receiving antipsychotic medication (2 first-generation, 43 second-generation, and 15 both). The Brief Psychiatric Rating Scale (BPRS), the Scale for the Assessment of Negative Symptoms (SANS) and the Scale for the Assessment of Positive Symptoms (SAPS) were obtained to evaluate the symptom severity. HC were recruited by advertisement. All HC have no current diagnosis of axis 1 or 2 disorders, substance dependence or abuse, or family history of psychosis. All subjects are right-handed with normal sight and color perception. Two groups of subjects were matched in age and educational level.

### Stimuli and Task

The experiment was run on the platform of Matlab 8.1 and Psychtoolbox 3. Subjects were seated at a distance of 50 cm away from an LCD monitor.

Each trial started with a fixation cross presented at the center of the screen, lasting for the time randomly chosen from [300, 350, 400, 450, 500 ms]. A set of colored shapes (squares and/or circles) were then shown on the screen on an invisible circle with 4° radius for 500 ms. Four conditions were used in this experiment: target size 1 / 3 × distractor size 0 / 2. Half of the subjects were instructed before the experiment started to remember colored squares (target items) and ignore colored circles (distractors) for the whole experiment and vice versa. Colored squares were 1.5° × 1.5° of visual angles and colored circles were 1.5° of visual angles in diameters. The sample array was shown for 500 ms, followed by a 900 ms delay period with only the fixation cross on the screen for memory retention. Then, an equal number of outlined shapes were presented at the same locations of the items shown in the sample array. One of the outlined shapes was bolded, indicating the target item at this cued location is to be recalled. Meanwhile, a randomly rotated color wheel was shown on the screen, with the inner and outer radius as 7.8° and 9.8° respectively. Subjects were instructed to recall and report the color of the bolded item by clicking on the color wheel using a computer mouse. Precise recall of the color was desired and the response time was unlimited. The 180 colors used in this experiment were selected from a circle (centered at L = 70, a = 20, b = 38, radius of 60) deriving from the CIE L*a*b color space. All subjects finished one block of 80 trials for each condition. The order of conditions was counterbalanced across subjects.

### Data analysis

The data with no distractor has been presented in reference (32)^32^. Comprehensive analysis of the distraction effect in this paper is new.

<u>Variable precision model</u>. The variable precision (VP) model was initially proposed by van den Berg *et al*^31,33^. The VP model proposes that the mean VWM resource levels declines as the target size assigned to individual items are not only continuous but also variable across items and trials. This variability in resource assignment results in trial-by-trial response errors. Moreover, the VP model also explicitly isolated the variability of behavior choice (e.g., motor or decision noise), which was ignored by most previous models in VWM.

For each item, the memory resources recruited *J* is defined as Fisher information 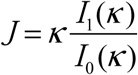, where *I* and *I*_0_ and *I*_1_ are modified Bessel functions of the first kind of order 0 and 1 respectively, with the concentration parameter *κ*. In the VP model, because *J* varies across items and trials, it is further assumed to follow a Gamma distribution with a mean of 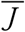 and scale parameter *τ*. Moreover, since the mean VWM resource decreases with target size *N* (Fig. 3A), we assume that the relationship between 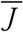 and *N* can be written in a power-law fashion 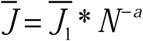, where 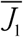 is the initial resources when only 1 item (*N* = 1) should be remembered in VWM and *α* is the decay exponent.

**Figure 2.**
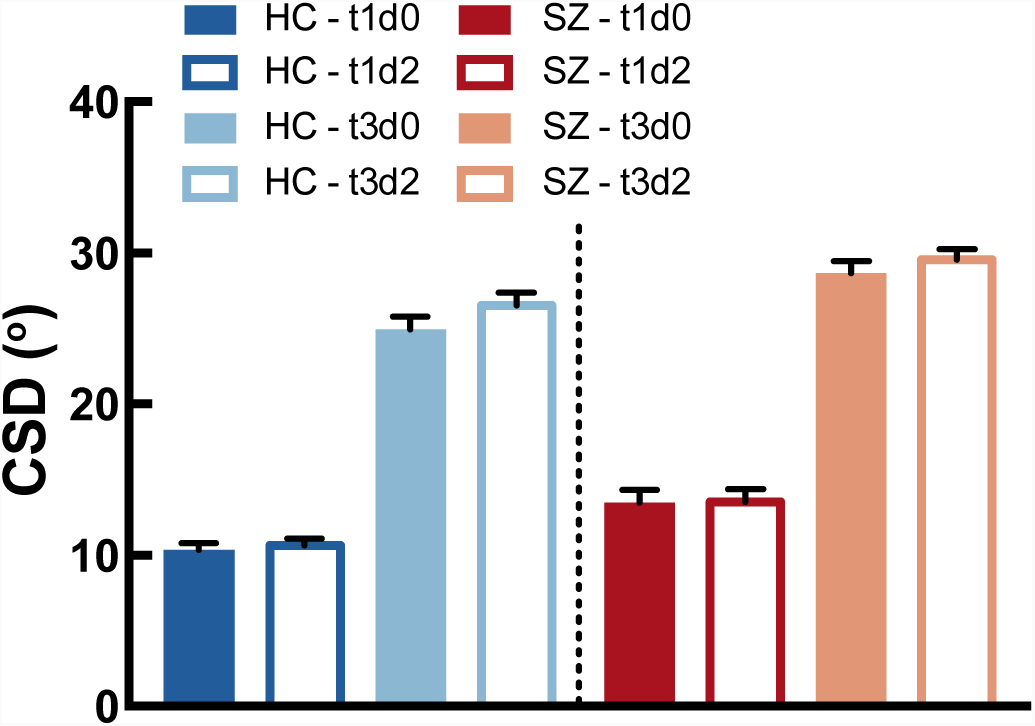
General memory load and distraction effects on both groups. A higher CSD indicates worse performance. Increasing the memory load and the distractor level worsen performance in both groups. Also, SZ showed generally worse VWM performance than HC. Moreover, distractors only impact VWM performance at high memory load (target size = 3). Error bars represent ±SEM across subjects. The letter “t” in the legend means “target size” and “d” means “distractor size”. For example, “t1d0” indicates target size = 1 and distractor size = 0.

**Figure 3.**
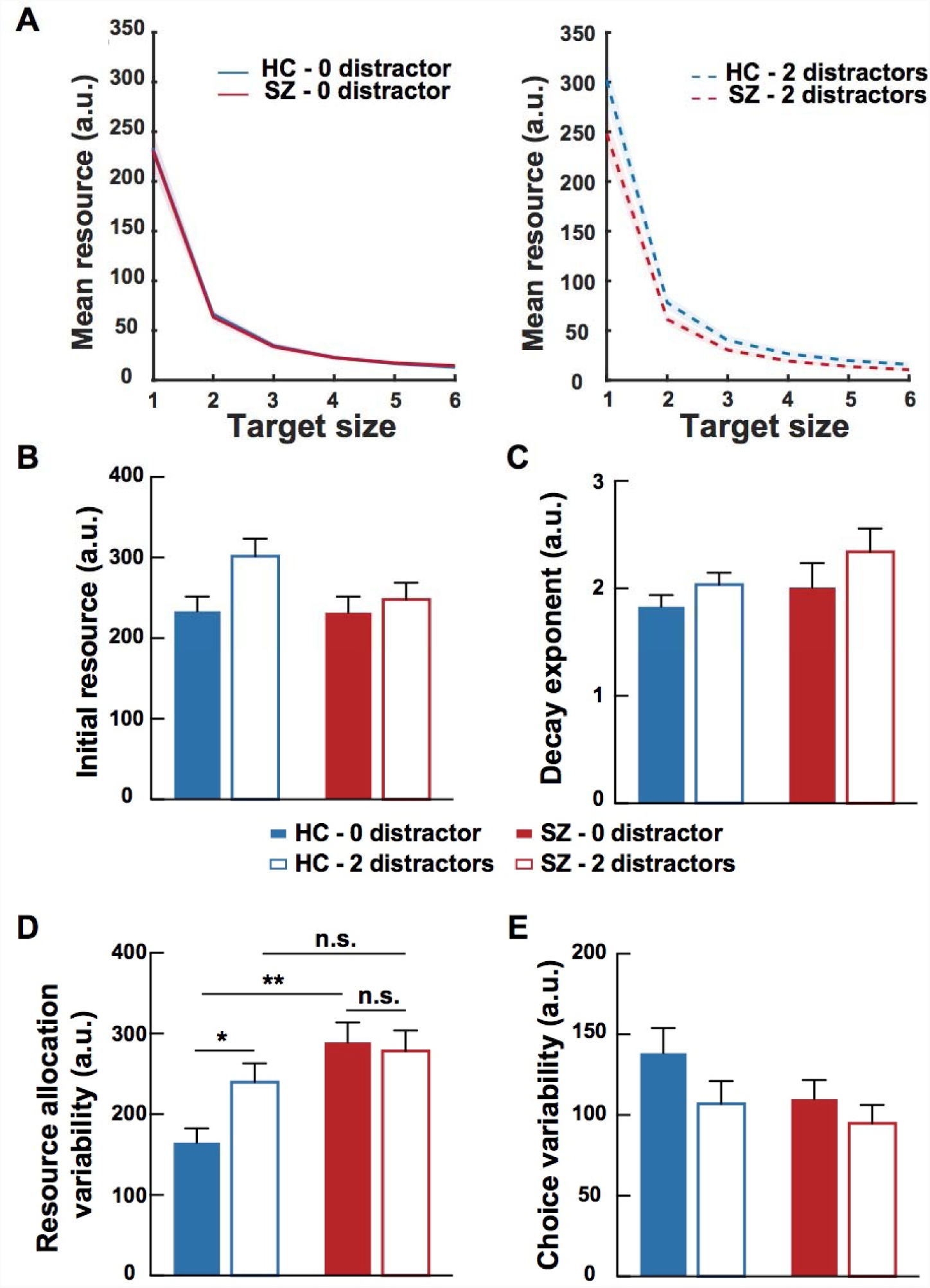
Effects of group, target size and distractor size on the four fitted parameters of the VP model. Panel A illustrates the mean resources as a function of the target size, which are generated by fitted initial resource (panel B) and decay exponent (panel C) values. Panels D and E illustrate the fitted resource allocation variability and choice variability respectively. The main group effect was only found in resource allocation variability (panel D). Precisely, SZ showed overall larger resource allocation variability than HC and adding distractors only elevated the resource allocation variability in HC but not SZ, indicating that SZ have stronger resilience to distraction than HC. No group × distractor size interaction was observed in initial resource, decay exponent and choice variability. Shaded areas in panel A and error bars in panels B to E denote ±SEM across subjects. Significance symbol conventions are *: p < 0.05; **: p < 0.01; n.s.: non-significant.

The model also assumes that the subject’ internal representations of stimuli are noisy and follow a von Mises distribution. Thus, the distribution of sensory measurement (*m*) given the input stimulus (*s*) can be written as:

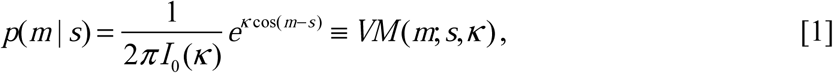

and we further assumes that subjects’ reported color (ŝ) shat also follows a von Mises distribution with the choice variability *K*_*r*_:

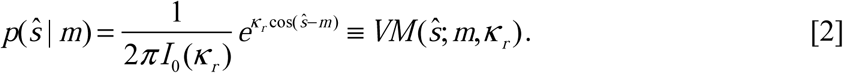

Taken together, there are four free parameters: 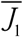, *α, τ* and *K*_*r*_ in the VP model.

### Model fitting

We fit the model separately for each subject. Because *J* is a variable across items and trials, we sampled it for 10000 times from the Gamma distribution with mean 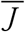 and scale parameter *τ*. We then used all these samples to calculate response probability in each trial.

We used the BADS optimization toolbox in MATLAB to search the best fitting parameters that maximize the likelihood of responses. To avoid the issue of local minima, we did the optimization process for 20 times with 20 different initial seeds. The parameters with the maximum likelihood were used as the best fitting parameters for a subject and were further used in the statistical process.

## RESULTS

### SZ make larger recall errors than HC

We set four experimental conditions (target size 1/3 x distractor size 0/3) for each group. In the modified color delay-estimation task, performance in a trial, denoted as “response error”, was defined as the distance between the true color and the reported color of the cued item in the circular color space. For each subject, circular standard deviations (CSD) of response errors in each experimental condition were calculated separately as indexes of VWM performance.

A 2 × 2 × 2 ANOVA was performed with the CSDs as the dependent variable (Fig. 2), target size (1/3) and distractor size (0/3) as the within-subject variables, and group (SZ/HC) as the between-subject variable. We observed the main effects of target size (*F(1,119)* = 935.650, *p* < 0.001, partial *η*^2^ = 0.887) and distractor size (*F(1,119)* = 8.909, *p* = 0.003, partial *η*^2^ = 0.070), indicating that behavioral performance in both groups declined as the memory load and the distraction level increased. These results also suggest that our experimental manipulation successfully induced the classical load effect and the distraction effect. A group difference was also found (*F(1,119)* = 12.716, *p* < 0.001, partial *η*^2^ = 0.097) and we confirmed a general worse VWM performance of SZ than HC, a result consistent with many previous studies showing the VWM deficits in schizophrenia^3–6^. We also found a significant interaction between target size and distractor size (*F(1,119)* = 4.486, *p* = 0.036, partial *η*^2^ = 0.036). Post hoc analysis showed that the distractors worsened VWM performance (*p* = 0.004) in the high target size (i.e., target size = 3) condition, whereas no distraction effect was detected in the low target size (i.e., target size = 1) condition (*p* = 1.000).

The key question we asked here was whether the distractors selectively impaired VWM processing in SZ. If yes, we should expect an interaction effect between distractor size and group as adding distractors might impose stronger performance deteriorations in SZ compared with HC. However, we did not find such interaction effect (*F(1,119)* = 0.820, *p* = 0.367, partial *η*^2^ = 0.007), indicating that adding distractors worsened performance in both groups and such distraction effect was not specific to SZ. Moreover, previous studies have suggested that distractibility deficits in SZ might be more prominent when the task becomes more challenging (e.g., higher memory load). However, no other significant interaction effect was noted with respect to the group variable (target size × group, *F(1,119)* = 0.139, *p* = 0.710, partial *η*^2^ = 0.001; target size × distractor size × group (*F(1,119)* = 0.137, *p* = 0.712, partial *η*^2^ = 0.001). These results were consistent with the previous studies^25,26,34^ showing that SZ exhibit generally worse VWM performance than HC but the memory load and distraction effect manifest similarly in both groups.

### Distractors elevate resource allocation variability in HC but not in SZ

Above analyses only focused on CSD—a summary statistics describing the variance of recall error distributions in each experimental condition. To further scrutinize the data, we employed the VP model (see Methods)—a Bayesian observer model describing the generative process of a behavioral choice in the delay-estimation task. The VP model has two major strengths. First, unlike the CSD as a summary statistical variable, the VP model is a probabilistic model that can utilize the data in every trial without losing any information. Second and more importantly, the VP model explicitly defines some key VWM components and characterizes the full generative process of the VWM task. Therefore, we can quantify the distraction effect on these VWM components.

We elaborated the details of the VP model here. First, the VP model estimates the initial resources when only one target is present. Second, the memory resources decline as a power function of target size and this decreasing trend can be described by the decay exponent. Third, the power function only specifies the mean resource at each target size level. The actual resources assigned to each item vary and follow a Gamma distribution with the variance as resource allocation variability. The amount of resources assigned to each item determines the precision of sensory measurement (i.e., memory representation) of the item. Forth, given the noisy representation, there exists choice variability describing the uncertainty from internal sensory representation to the outcome behavioral choice. We estimated the four parameters (i.e., initial resources, decay exponent, resource allocation variability and choice variability) on each subject and separately on two distractor size levels.

We performed a 2 × 2 ANOVA with distractor size as the within-subject variable, group as the between-subject variable, and the four estimated parameters of the VP model as the dependent variables. We observed a main effect of group in resource allocation variability (*F(1,119)* = 9.863, *p* = 0.002, partial *η*^2^ = 0.077), showing an overall higher resource allocation variability in SZ compared to HC (Fig. 3D). This result is consistent with our earlier work^32^. The main effect of group was not significant in the other three parameters. Particularly, we did not observe a significant main effect of initial resource and decay exponent, two factors that control the amount of memory resources. Intuitively, these results suggest that SZ might have the same amount of memory resources, but they distributed the resources across targets in a very heterogeneous manner.

We also found a main effect of distractor size on initial resource (*F(1,119)* = 5.559, *p* = 0.020, partial *η*^2^ = 0.045) and a marginal significant main effects on decay exponent (*F(1,119)* = 3.882, *p* = 0.051, partial *η*^2^ = 0.032). We speculate that adding distractors greatly enhanced the task difficulty and consequently forced subjects to internally utilize more resources to memorize targets. There were no main effects of distractor size on choice variability (*F(1,119)* = 3.528, *p* = 0.063, partial *η*^2^ = 0.029) and resource allocation variability (*F(1,119)* = 2.862, *p* = 0.093, partial *η*^2^ = 0.023). Note that these main effects manifest in both groups not specific for SZ.

More importantly, to examine the distraction effect, the key is to examine the interaction effect between group and distractor size. If SZ have deficits in distractibility, we should expect that adding distractors imposes significantly larger interferences on VWM processing in SZ but compared with HC. We indeed observed a significant interaction effect between group and distractor size (F(1,119) = 5.062, p = 0.026, partial *η*^2^ = 0.041) (Fig. 3D) in resource allocation variability. However, post hoc analysis suggested that adding distractors only increased the resource allocation variability in HC (*p* = 0.036) but had little impact on SZ (*p* = 0.999). This is surprising since elevated distractibility has long been proposed as a core executive function deficit in SZ. On the contrary, we found a more prominent distraction effect in HC rather in SZ, indicating a relatively higher resilience to distraction in SZ. We did not find such interaction effect in all other three parameters (initial resource, *F(1,119)* = 2.042, *p* = 0.156, partial *η*^2^ = 0.017; decay exponent, *F(1,119)* = 0.236, *p* = 0.628, partial *η*^2^ = 0.002; choice variability, *F(1,119)* = 0.430, *p* = 0.513, partial *η*^2^ = 0.004).

These results also suggest the critical role of resource allocation variability since we did not find the interaction effect of group and distractor size, as well as their interaction on other three VP model parameters (see full statistical results in Supplementary Materials note 1). Resource allocation variability is a relatively new concept in VWM and has increasingly been regarded as one of the key determinants for VWM performance^31^. Also, our earlier work confirmed its contribution to schizophrenic pathology^32^. Recent studies have shown that it is not only a key component in VWM but might be also a very general property in sensory processing^35^.

## DISCUSSION

Visual working memory and distractibility have long been recognized as core executive functions. Despite the widely documented behavioral deficits of SZ in these two domains, little is known with respect to the computational mechanisms underlying these deficits. This arises from two major obstacles: (1) few studies have attempted to integrate two cognitive functions within the same experimental paradigm; (2) the computational models that describe the internal processes have been lacking. To circumvent these, we modified the classical VWM delay-estimation task to deliberately incorporate distractors and employed the VP model to distinguish several VWM key components. We set two distractor conditions (distractor size 0/3) and used the VP model to estimate the VWM components separately under these two conditions. We made two major observations: (1) the variability of allocation memory resources was generally larger in SZ;(2) adding distractors enlarged the resource allocation variability in HC but had little impact on that in SZ. These results highlight the significance of resource allocation variability in mediating VWM performance and demonstrate an unexpected higher resilience to distraction during VWM in SZ.

The finding of enhanced resource allocation variability is of unique significance for understanding VWM deficits in SZ. This finding has been systematically evaluated in our prior work^32^. In that study, we compared several influential models in VWM literature and compare results between SZ and HC. We found that the only difference between the two groups lies in resource allocation variability not the amount of memory resources. This result suggests that SZ have the same amount of mean resources as HC at each target size level, but the resources assigned to individual items exhibit larger variability around this mean value. For example, assume that, given three targets, both SZ and HC have *r* units of mean resource across three targets. But the actual resources assigned to each item vary around this mean value (i.e., *r*+0.1, *r*-0.2). SZ exhibit overall larger variability (e.g., *r*+3, r-2) than HC (e.g., *r*+0.3, *r*-0.2). Note that this mechanism is fundamentally different from elevated attentional lapse or general deficits in filtering distraction. Elevated attentional lapse will lead to more guessing trials and the general deficits in filtering distraction will allow more resources assigned to distractors. Therefore, these mechanisms predict that the mean resources will be overall reduced in SZ. However, we did not observe the significant group differences in memory resources (Fig. 3A).

The unexpected enhanced resilience to distraction in resource allocation variability provides a new perspective for understanding distractibility in SZ. We confirmed that behavioral performance of SZ is in general worse than HC, a well-established finding in many previous studies^3–6^. However, in the analyses of behavioral performance, we indeed observe significant effects of memory load and distraction but both effects manifest similarly in both groups. There was no stronger distraction effect specific for SZ. Most previous studies employed a similar approach and only focused on behavioral performance. We made a further stride here and examined the distraction effect on individual VWM computational components. Results showed that adding distractors only significantly raise the resource allocation variability in HC but not in SZ. This is the key contribution of our work. Our approach allows us to provide a deeper mechanistic interpretation rather than only reporting the quantitative behavioral deficits in SZ. Note that our approach here is to fit the VP model separately to the data at two distraction conditions and then examine the differences in the estimated parameters. An alternative approach is to directly incorporate the distraction effect into the generative process, which has been recently pursued in Ni & Ma^36^ and Shen & Ma^35^. The latter approach permits to compare different computational models so as to ground different theories. Future work might continue to explore this line of research.

At first glance, higher resilience to distraction in the VWM resource allocation suggests a cognitive advantage in SZ. However, this might also imply less flexible cognitive control in SZ. For example, there has been shown that SZ tend to allocate their VWM resources more intensely and narrowly than HC^37^, a phenomenon called “hyperfocusing”. If SZ distribute too many resources on a small set of visual objects, they may have trouble in flexibly switching to new objects. Hyperfocusing might be particularly problematic in VWM tasks since one of the key features of VWM is to flexibly and dynamically maintain representations of multiple objects. The hyperfocusing mechanism might explain both the elevated resource allocation variability and higher distraction resilience in SZ. In our task, hyperfocusing on a subset of targets avoids the interference of distractor. Again, note that the “side effect” of hyperfocusing might be the lack of ability to flexibly switch to different sources of information ^38^. Also, the atypical ability in task switching has also been discovered in other special populations, such as aging^39,40^, ADHD^41^.

What are the neural mechanisms underlying VWM deficits and distraction effects in SZ? A recent study has identified the superior intraparietal sulcus (IPS) as the cortical region controlling resource allocation variability ^42^. SZ patients have also been found the atypical neural processing in this region^43^. On the other hand, the distraction effect on neural processing has been broadly found in attention and cognitive control networks^44^. Especially, SZ exhibited abnormal neural processing when distractors are present and cortical activity in high-level brain regions (i.e., dorsolateral prefrontal cortex) is correlated with negative symptoms^45^. However, no study has combined the VWM and distractors paradigm and measured neural activity in SZ. Also, it is unclear how other computational components of VWM are implemented in the brain. Future studies might need to combine computational modeling, neural measurements and behavioral testing to systematically address this issue.

Taken together, in this study we combined the standard VWM and distractor paradigms to examine the distraction effect during VWM in both SZ and HC. We replicated the standard memory load and distraction effects in both groups. We also found general worse VWM performance in SZ. But we did not observe a significant higher distraction effect in SZ. Further modeling analyses revealed that distractors elevate resource allocation variability during VWM in HC but not in SZ. This unexpected higher resilience to distraction in SZ provides new evidence for the cognitive deficits of SZ. Such unexpected higher resilience and less flexible cognitive control might be two sides of the same coin.

## ACKNOWLEDGMENTS

This work was supported by the National Social Science Foundation of China (17ZDA323), the National Key Fundamental Research Program of China (2013CB329501), the Major Program of Science and Technology Commission Shanghai Municipal (17JC1404100), the Fundamental Research Funds for the Central Universities (2018ECNU-QKT015), and the NYU-ECNU Institute of Brain and Cognitive Science at NYU (YK).

## AUTHOR CONTRIBUTIONS

YZ, RZ, & YK developed research idea and study concepts; YZ & YK design the experiment; XR and LZ collected the data; RZ & YZ performed the data analyses and modeling; RZ&YZ wrote the manuscript.

